# P-TEFb activation by RBM7 shapes a pro-survival transcriptional response to genotoxic stress

**DOI:** 10.1101/394239

**Authors:** Andrii Bugai, Alexandre J.C. Quaresma, Caroline C. Friedel, Tina Lenasi, Christopher R. Sibley, Petra Kukanja, Koh Fujinaga, Melanie Blasius, Thomas Hennig, Jernej Ule, Lars Dölken, Matjaz Barboric

## Abstract

Cellular DNA damage response (DDR) involves dramatic transcriptional alterations, the mechanisms of which remain ill-defined. Given the centrality of RNA polymerase II (Pol II) promoter-proximal pause release in transcriptional control, we evaluated its importance in DDR. Here we show that following genotoxic stress, the RNA-binding motif protein 7 (RBM7) stimulates Pol II elongation and promotes cell viability by activating the positive transcription elongation factor b (P-TEFb). This is mediated by genotoxic stress-enhanced binding of RBM7 to 7SK snRNA (7SK), the scaffold of the 7SK small nuclear ribonucleoprotein (7SK snRNP) which inhibits P-TEFb. In turn, P-TEFb relocates from 7SK snRNP to chromatin to induce transcription of short units including key DDR genes and multiple classes of non-coding RNAs. Critically, interfering with RBM7 or P-TEFb provokes cellular hypersensitivity to DNA damage-inducing agents through activation of apoptotic program. By alleviating the inhibition of P-TEFb, RBM7 thus facilitates Pol II elongation to enable a pro-survival transcriptional response that is crucial for cell fate upon genotoxic insult. Our work uncovers a new paradigm in stress-dependent control of Pol II pause release, and offers the promise for designing novel anti-cancer interventions using RBM7 and P-TEFb antagonists in combination with DNA-damaging chemotherapeutics.

## INTRODUCTION

Cellular DDR has evolved to detect and repair lesions that are generated continuously by external and internal DNA-damaging agents (Hoeijmakers, 2009). In parallel, activation of DDR halts progression of the cell-cycle to provide time for repair, of which outcome determines whether cells will re-enter the cell-cycle and continue with their physiological program or die by senescence or apoptosis. Defective DDR can lead to genomic instability underlying many diseases, including hematological disorders and cancer (Jackson and Bartek, 2009). Importantly, it is widely accepted that cells need to shut down Pol II transcription in response to ultraviolet light (UV)-induced bulky DNA lesions and other types of DNA damage, which can facilitate the repair and limit the production of abnormal transcripts (Giono et al., 2016). While transcription can be inhibited transiently at initiation and elongation stages (Awwad et al., 2017; Rockx et al., 2000; Williamson et al., 2017), or irreversibly through degradation of stalled Pol II (Wilson et al., 2013), it is eventually restored once the damage is corrected. However, mechanisms and significance of mounting transcriptional activation following genotoxic stress remain poorly understood.

Transition of Pol II from promoter-proximal pausing to productive elongation represents a critical rate-limiting step in metazoan gene expression (Jonkers and Lis, 2015; Zhou et al., 2012). At most active genes, Pol II transcribes 20-100 nucleotides from transcription start site before productive elongation is paused by the multi-subunit negative transcription elongation factors (N-TEFs), consisting of negative elongation factor (NELF) and DRB-sensitivity inducing factor (DSIF). The release of paused Pol II genome-wide is stimulated by P-TEFb kinase, which is composed of the catalytic CDK9 and a regulatory CycT1 or CycT2 subunit (Gressel et al., 2017; Jonkers et al., 2014; Mayer et al., 2015; Peterlin and Price, 2006). Upon its recruitment or activation at target gene promoters or enhancers, P-TEFb phosphorylates serine 2 residues (Ser2-P) within the C-terminal domain (CTD) Y^1^S^2^P^3^T^4^S^5^P^6^S^7^ heptapeptide repeats of the largest Pol II subunit RPB1, as well as NELF-E and the SPT5 subunit of DSIF, leading to productive Pol II elongation (Jonkers and Lis, 2015; McNamara et al., 2016; Quaresma et al., 2016; Zhou et al., 2012). In addition, clearance of Pol II from promoter-proximal pause sites enables new transcriptional initiation, augmenting the production of RNA (Gressel et al., 2017; Shao and Zeitlinger, 2017).

Considering that P-TEFb is crucial for prompt expression of stimulus-induced genes (Liu et al., 2015), we reasoned that it might feature prominently in stimulating Pol II transcription following DNA damage. Importantly, a major fraction of P-TEFb resides within 7SK snRNP, in which coordinated actions of the scaffolding 7SK and three canonical RNA binding proteins (RBPs) inhibit the kinase. While the 7SK g-methylphosphate capping enzyme (MePCE) and La-related protein 7 (LARP7) stabilize 7SK to form the core of 7SK snRNP, HEXIM1 or its paralogue HEXIM2 subsequently interacts with 7SK of the core to bind and inhibit P-TEFb (Guo and Price, 2013; Quaresma et al., 2016). Of note, activation of P-TEFb through its release from 7SK snRNP, as exemplified most prominently by the RNA binding HIV-1 transcriptional transactivator Tat, has emerged as a key regulatory event that stimulates Pol II pause release at specific genes (Quaresma et al., 2016). Given that cellular RBPs are emerging as important effectors in DDR (Dutertre et al., 2014; Wickramasinghe and Venkitaraman, 2016) and transcription (Quaresma et al., 2016; Skalska et al., 2017), we reasoned that a protein from this class could mediate genotoxic stress-induced P-TEFb activation and thus Pol II transcription via 7SK snRNP.

In this study, we focused our efforts on the ubiquitously expressed RBM7, which promotes survival of cells following DNA damage generated by UV or its mimicking genotoxic and carcinogenic chemical 4-nitroquinoline 1-oxide (4-NQO) (Blasius et al., 2014). RBM7 binds the DExH/D box RNA helicase hMTR4 (also known as SKIV2L2) through the bridging zinc knuckle protein ZCCHC8, forming the nuclear exosome targeting complex (NEXT) (Falk et al., 2016; Lubas et al., 2011). Through the RNA-binding capacity of RBM7, NEXT promotes degradation of multiple RNA classes, including upstream antisense (uaRNAs) and enhancer-derived (eRNAs) transcripts, as well as 3′ end extended forms of snRNAs, snoRNAs and histone gene transcripts (Andersen et al., 2013; Hallais et al., 2013; Hrossova et al., 2015; Lubas et al., 2011). Furthermore, NEXT targets pre-mRNAs for decay and/or processing of intron-embedded snoRNAs and miRNAs (Lubas et al., 2015), possibly via the interaction between RBM7 and spliceosomal SF3b complex (Falk et al., 2016). Finally, NEXT associates with the cap-binding protein complex and functions in connecting transcription termination with exosomal degradation (Andersen et al., 2013; Hallais et al., 2013). Although the linkage between RBM7 and DDR has been reported (Blasius et al., 2014), it remains unclear how does RBM7 exert its pro-survival function. Here we uncovered an unexpected interplay between RBM7 and 7SK snRNP in the wake of DNA damage, shining a spotlight on the central role of P-TEFb kinase in shaping a transcriptional response that is crucial for viability of genotoxic-stressed cells.

## RESULTS

### iCLIP reveals DNA damage-enhanced interaction between RBM7 and 7SK

The vital role of RBM7 in DDR led us to postulate that defining the RNA interactome of RBM7 under unchallenged and DNA damage conditions should disclose insights into the mechanisms underlying its critical function in genotoxic-stressed cells. Thus, we performed individual-nucleotide resolution UV crosslinking and immunoprecipitation (iCLIP) assay in human HEK 293 Flp-In T-Rex (HEK 293) cells that expressed 3X-FLAG epitope-tagged RBM7 (F-RBM7). Because UV irradiation yields RNA-protein crosslinks immediately and thus prior to DDR activation, we instead employed its mimetic 4-NQO. Notably, 4-NQO metabolite 4-hydroxyaminoquinolone 1-oxide forms bulky DNA adducts on purines, which are removed by nucleotide excision repair (Ikenaga et al., 1977; Tada and Tada, 1976). Consistent with the previous report (Lubas et al., 2015), RBM7 bound a diverse set of RNAs, including genic, intergenic and non-coding (nc) RNAs (Figures 1A and S1). We ranked these RNAs by the change in binding following 4-NQO exposure, which showed increased RBM7 binding to snRNAs, including 7SK, spliceosomal snRNAs, and other ncRNAs, and decreased RBM7 binding to specific pre-mRNAs (Tables S1 and S2). As expected, 4-NQO exposure led to a dramatic increase in *γ*-H2AX foci formation, confirming activation of DDR (Figure S2A). Given the pivotal role of 7SK snRNP in regulating Pol II pause release (McNamara et al., 2016; Quaresma et al., 2016), we set out to investigate the potential connection between RBM7 and 7SK in controlling Pol II transcription following genotoxic stress.

**Figure 1.**
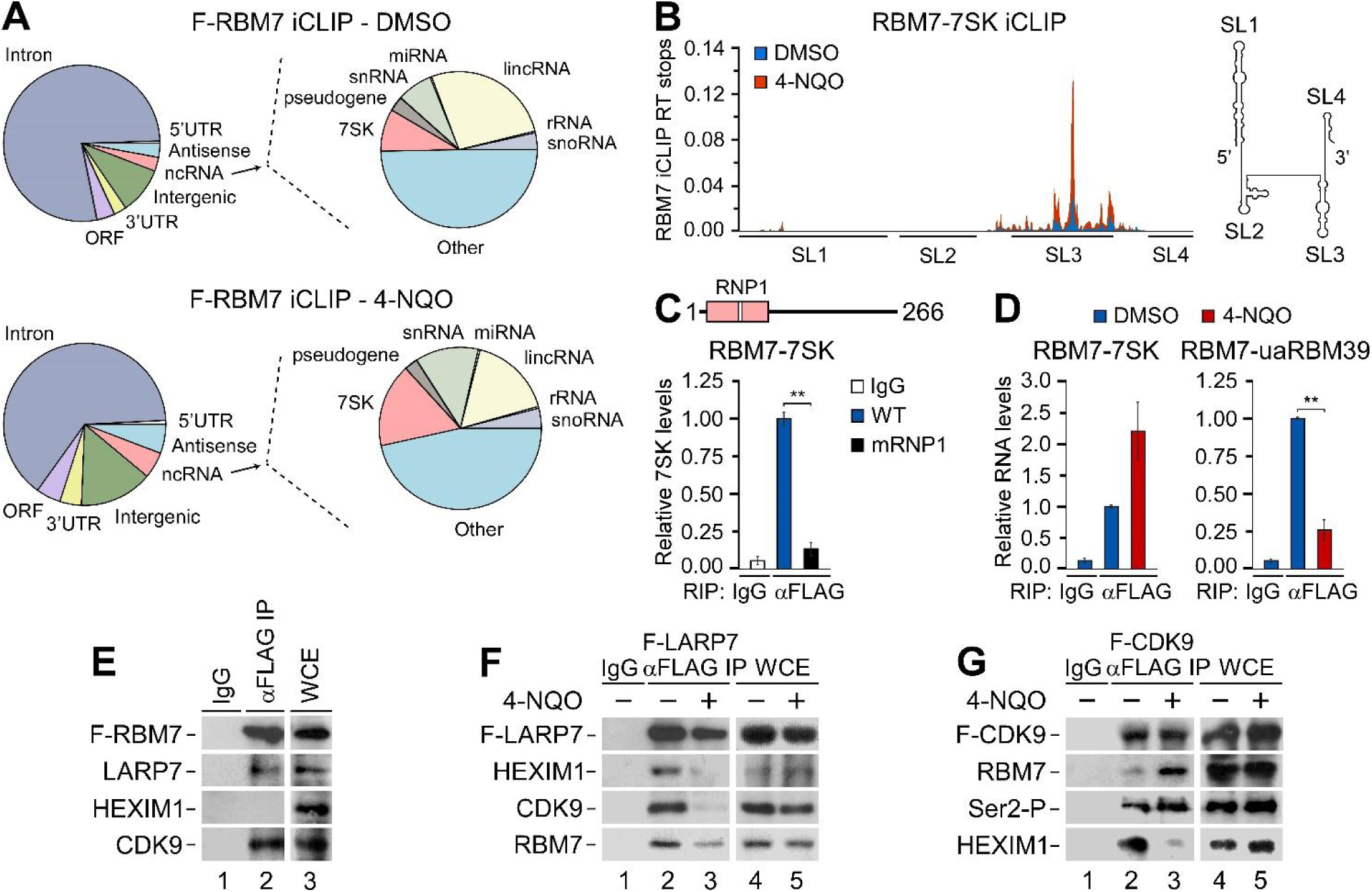
DNA damage induces the interaction of RBM7 with 7SK and relocation of P-TEFb from 7SK snRNP to Pol II. (A) Distribution charts of unique tags derived from the mock and 4-NQO treated RBM7 libraries based on percentages of the total iCLIP reads and mapped to the indicated RNA classes. Charts on the right show distribution of the indicated types of ncRNA. (B) 3X-FLAG epitope-tagged RBM7 protein (F-RBM7) iCLIP reads mapped to 7SK. Positions of the four stem–loops (SL1–4) are indicated below the iCLIP reads and on a 7SK secondary structure model. (C) Quantitative RNA immunoprecipitation (RIP-qPCR) of 7SK in WT and mRNP1 F-RBM7 immuno-purifications from HEK 293 whole-cell extracts (WCE). RBM7 with RRM (in pink) and the position of RNP1 (white stripe) is depicted on top. (D) RIP-qPCR of 7SK and uaRBM39 in F-RBM7 immuno-purifications from HEK 293 WCE. Conditions with (red bars) and without (blue bars) 4-NQO are shown. Results in C and D are presented as the mean ± s.e.m. (n = 3 independent experiments). **, P < 0.01. (E,F) Co-immunoprecipitation of F-RBM7 and 3X-FLAG epitope-tagged LARP7 protein (F-LARP7) with 7SK snRNP from HEK 293 WCE. Conditions with (+) and without (-) 4-NQO are shown. (G) Co-immunoprecipitation of 3X-FLAG epitope-tagged CDK9 protein (F-CDK9) with the indicated proteins from HEK 293 WCE. Conditions with (+) and without (-) 4-NQO are shown.

We next plotted the RBM7 iCLIP reads on the secondary structure model of 7SK. This revealed that RBM7 binds selectively stem-loop 3 (SL3) of 7SK (Figure 1B), thereby placing it between MePCE/HEXIM1 and LARP7 that bind the SL1 and SL4, respectively. Quantitative RNA immunoprecipitation (RIP-qPCR) assays proved the iCLIP result to be specific since substituting the conserved Lys60, Phe62 and Phe64 residues of the ribonucleoprotein 1 (RNP1) motif within the RNA recognition motif (RRM) of RBM7 to alanines (mRNP1) abrogated the interaction (Figure 1C). Furthermore, we confirmed that 4-NQO exposure increased the binding of RBM7 with 7SK, while it decreased its binding with the uaRNA of RBM39 (Figure 1D), which is consistent with the DNA damage-induced release of uaRNAs from NEXT (Blasius et al., 2014; Tiedje et al., 2015). These results identify RBM7 as a novel 7SK-interacting protein and suggest a possible role of RBM7 in facilitating the cellular response to genotoxic stress via 7SK snRNP.

### DNA damage stimulates the relocation of P-TEFb and RBM7 from 7SK snRNP to Pol II

To test if RBM7 binds the 7SK snRNP, we performed a co-immunoprecipitation (co-IP) analysis in HEK 293 cells. This revealed the interaction of F-RBM7 with endogenous LARP7 and CDK9, but not HEXIM1 (Figure 1E). Importantly, 4-NQO treatment triggered the release of endogenous CDK9, HEXIM1 and RBM7 from the 7SK snRNP in HEK 293 and HeLa Flp-In (HeLa) cells, as determined by co-IP analysis with 3X-FLAG epitope-tagged LARP7 (F-LARP7) as the bait and by glycerol gradient centrifugation analysis, respectively (Figures 1F and S2B). Likewise, exposing HeLa cells to UV and primary human foreskin fibroblasts (HFF-1) to 4-NQO released CDK9 from HEXIM1 (Figures S2C and S2D).

To follow the relocation of P-TEFb upon genotoxic stress further, we examined its interaction with RBM7 and Pol II by conducting a series of co-IP and bimolecular fluorescence complementation (BiFC) assays. 4-NQO treatment enhanced the interaction of 3X-FLAG epitope-tagged CDK9 (F-CDK9) with RBM7 (Figure 1G), suggesting that the initial enhanced binding between RBM7 and 7SK is followed by the release of RBM7 in complex with P-TEFb. Notably, interaction of the released F-CDK9 with the transcriptionally engaged Ser2-P form of Pol II increased following DNA damage (Figures 1G). Likewise, 4-NQO treatment enhanced the binding between F-CDK9 and total Pol II (Figure S3A). To extend these findings, we monitored the interaction between P-TEFb and the CTD of Pol II in living HeLa cells using the visualization of P-TEFb activation assay (Fujinaga et al., 2015). In this approach, BiFC of transiently expressed YC-P-TEFb and YN-CTD chimera containing the C- and N-terminal region of yellow fluorescent protein (YFP), respectively, indicates the interaction (Figure S3B). 4-NQO exposure increased the number of YFP-positive cells to a similar extent as the known P-TEFb releasing agents SAHA and JQ1 (Bartholomeeusen et al., 2012; Contreras et al., 2009), which corroborated our co-IP results (Figures S3C and S3D). These findings establish that RBM7 binds 7SK snRNP, and that upon genotoxic stress, P-TEFb relocates to the CTD of Pol II.

### RBM7 releases P-TEFb from the core of 7SK snRNP upon DNA damage

Based on the observations gathered thus far, we hypothesized that following DNA damage, RBM7 activates P-TEFb by promoting its release from the 7SK snRNP. As part of NEXT, RBM7 could recruit the exosome to 7SK snRNP, resulting in its disintegration via 7SK nucleolysis. However, as determined by co-IP and RIP-qPCR experiments, the 4-NQO-induced release of CDK9 and HEXIM1 from F-LARP7 left the core of 7SK snRNP intact (Figure 2A). Namely, F-LARP7 and MePCE remained bound during DDR (Figure 2A), and neither the total nor the LARP7-bound levels of 7SK were altered considerably (Figure 2B). Confirming integrity of the core, we detected its increased association with hnRNP1 A1 (Figure 2A), which binds 7SK as it replaces P-TEFb and HEXIM1 during transcriptional inhibition (Quaresma et al., 2016; Zhou et al., 2012). Conversely, the levels of HEXIM1-bound 7SK decreased as anticipated (Figure 2B). Unlike the reconstitution of 7SK snRNP upon removing a P-TEFb inhibitor flavopiridol (FP) from cell culture medium, the washout of 4-NQO did not induce re-sequestration of CDK9 and HEXIM1 into the 7SK snRNP, most likely due to the remaining DNA damage as suggested by the persisting levels of phosphorylated *γ*-H2AX (Figures 2A and S4). Thus, RBM7 does not employ the RNA exosome to activate P-TEFb.

**Figure 2.**
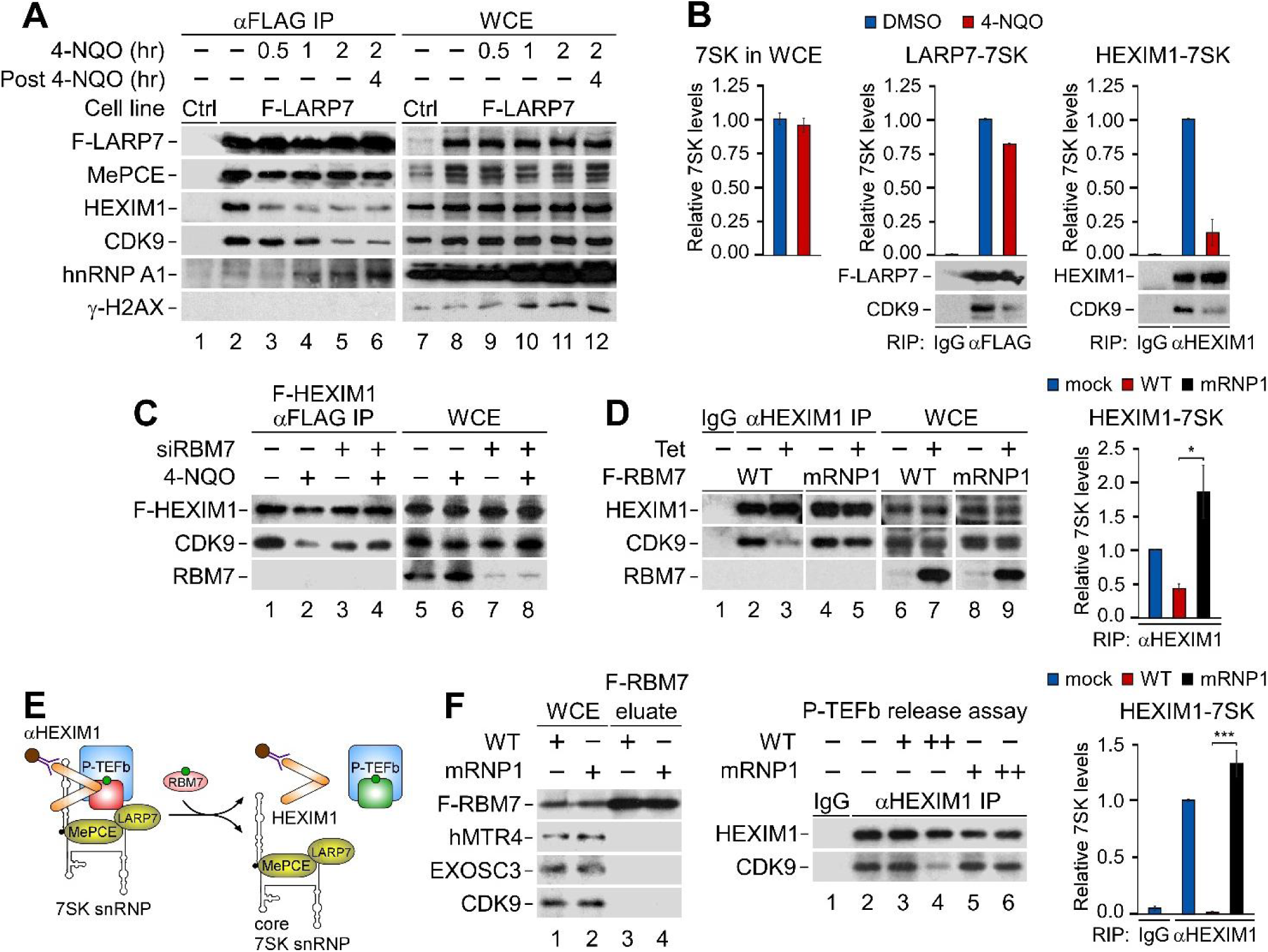
RBM7 releases P-TEFb from the core of 7SK snRNP upon DNA damage. (A) Co-immunoprecipitation of 3X-FLAG epitope-tagged LARP7 protein (F-LARP7) with 7SK snRNP and *γ*-H2AX from HEK 293 WCE. Conditions with (in hours) and without (-) 4-NQO are shown. (B) RT-qPCR of 7SK (left) and RIP-qPCR of 7SK in F-LARP7 (middle) and HEXIM1 (right) immuno-purifications (IP) from HEK 293 WCE. Conditions with (red bars) and without (blue bars) 4-NQO are shown. Results are presented as the mean ± s.e.m. (n = 2 independent experiments). Protein levels in the IP are shown below the graphs. (C) Co-immunoprecipitation of 3X-FLAG epitope-tagged HEXIM1 protein (F-HEXIM1) with CDK9 and RBM7 from HEK 293 WCE. Conditions with control (-) and RBM7 siRNA #1 (+), and with (+) and without (-) 4-NQO are shown. (D) (Left) Co-immunoprecipitation of HEXIM1 with CDK9 from HEK 293 WCE containing WT and mRNP1 F-RBM7. Conditions with (+) and without (-) F-RBM7 induction by tetracycline (Tet) are shown. (Right) RIP-qPCR of 7SK in HEXIM1 IP from HEK 293 WCE containing WT and mRNP1 F-RBM7. Conditions with WT (red bars), mRNP1 (black bars) and without (blue bars) F-RBM7 induction by tetracycline (Tet) are shown. Results are presented as the mean ± s.e.m. (n = 3 independent experiments). *, P < 0.05, determined by Student’s *t* test. (E) Cartoon depicting P-TEFb release assay. Release of inactive P-TEFb (CDK9 in red) from 7SK snRNP abrogates the interaction of HEXIM1 with P-TEFb, resulting in P-TEFb activation (CDK9 in green). (F) (Left) Eluates of immuno-purified WT and mRNP1 F-RBM7 from HEK 293 WCE. (Middle) Western blot analysis of P-TEFb release from HEXIM1 immuno-purified (αHEXIM1 IP) 7SK snRNP by the F-RBM7 proteins. Control (-, lanes 1 and 2) and conditions with (+) increasing amounts of WT and mRNP1 F-RBM7 are shown. (Right) RIP-qPCR of 7SK in HEXIM1 IP. Conditions with WT (red bars), mRNP1 (black bars) and without (blue bars) F-RBM7 incubation are shown. Results are presented as the mean ± s.e.m. (n = 4 independent experiments). ***, P < 0.001, determined by Student’s *t* test.

Alternatively, RBM7 could act on 7SK snRNP directly to release P-TEFb during DDR. To determine if RBM7 is critical for the release, we depleted it using RNA interference (RNAi). Indeed, the 4-NQO-induced release of CDK9 from 3X-FLAG epitope-tagged HEXIM1 (F-HEXIM) was abrogated in RBM7 knockdown cells (Figure 2C). In a complementary approach, inducible expression of F-RBM7 in HEK 293 cells decreased the interactions of endogenous HEXIM1 with CDK9 and 7SK, but this effect was diminished when using the 7SK binding-deficient mRNP1 F-RBM7 (Figure 2D). Finally, we examined if RBM7 released CDK9 from 7SK snRNP in vitro by using a P-TEFb release assay (Figure 2E). We immuno-purified the wild-type (WT) and mRNP1 F-RBM7 from HEK 293 cells under stringent conditions to strip-off its binding partners including subunits of NEXT, RNA exosome and P-TEFb (Figure 2F). Next, we incubated increasing amounts of the F-RBM7 proteins with the immobilized 7SK snRNP isolated from HeLa cells using an anti-HEXIM1 antibody. Confirming our prediction, F-RBM7 released CDK9 and 7SK from HEXIM1, while the mRNP1 F-RBM7 failed to do so (Figure 2F). Together, these results show that following genotoxic stress, RBM7 can release P-TEFb from the core of 7SK snRNP directly.

### P-TEFb directs transcriptional activation by Pol II in response to genotoxic stress

To disclose if active P-TEFb is vital for shaping a transcriptional response to genotoxic stress, we sequenced newly transcribed RNAs under unchallenged and DNA damage conditions. Because the onset of P-TEFb activation occurs within two hours of genotoxic stress (Figure 2A), we treated HeLa cells for one and two hours with 4-NQO. During the last 30 minutes, we labeled transcripts with the nucleoside analogue 4-thiouridine (4sU), which enables their isolation and sequencing (4sU-seq) (Figure 3A). Concurrently with 4-NQO, we also exposed the cells to a sub-optimal concentration of FP to evaluate if pharmacological inhibition of P-TEFb attenuates the response. Based on the reproducible 4sU-seq data (Figure S5), we determined differentially expressed (DE) mRNAs, long intergenic non-coding (linc) RNAs, uaRNAs and enhancer-derived eRNAs (p-value ≤ 0.01; expression fold-change ≥ 2; Tables S3-S6). This revealed that over a third of the DE coding genes and most of the DE ncRNAs are up-regulated (Figures 3A and 3B). Importantly, the stimulated Pol II transcription, which was similar at both durations of 4-NQO treatments (r=0.81 for eRNAs to 0.89 for uaRNAs; Figure S6), was highly dependent on active P-TEFb (Figure 3B). Notably, the response to 4-NQO was rather specific as only 10-13% of mRNAs and uaRNAs, and 3-5% of lincRNAs and eRNAs underwent expression changes (Table S7). In accordance with the previous work on transcriptional response to UV (McKay et al., 2004; Williamson et al., 2017), the up-regulated coding genes were dramatically shorter than the down-regulated genes (Figures 3C and S7). Finally, p53, bulky DNA adduct-inducers camptothecin and cisplatin, as well as DNA-damaging doxorubicin and sirolimus were identified by Ingenuity Pathway Analysis (IPA) Upstream Regulator analytic as top regulators that could yield the 4-NQO-like response at coding genes (Table S8), suggesting that DNA damage had indeed mediated the transcriptional changes.

**Figure 3.**
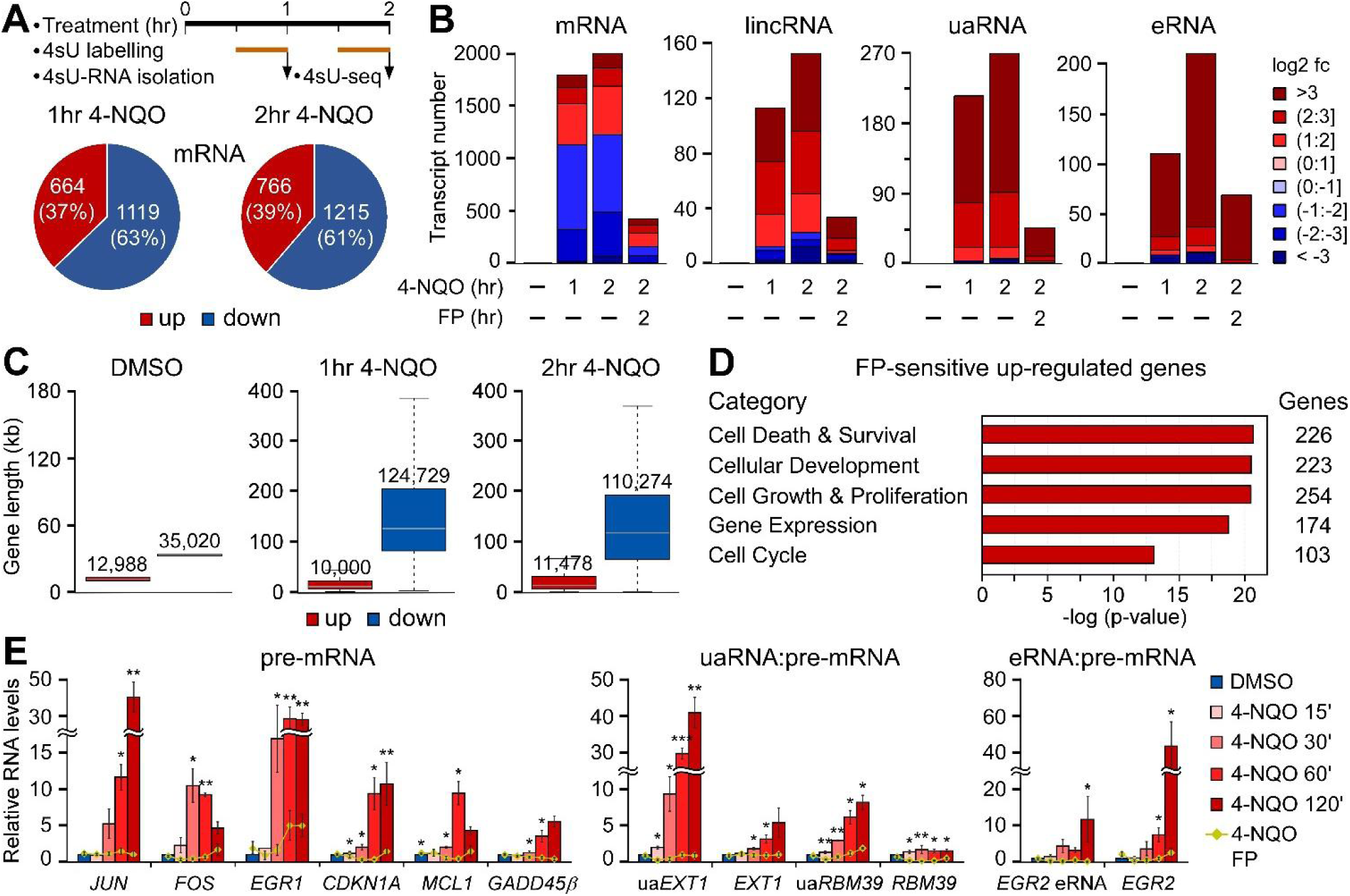
Active P-TEFb is vital for Pol II transcriptional response to DNA damage. (A) (Top) Schematic depicting major steps in the generation of 4sU-labeled transcripts (4sU RNA) for 4sU-seq. (Bottom) Pie charts showing the fractions of DE protein-coding genes (mRNA) upon one and two hours of 4-NQO treatment of HeLa cells as assessed by 4sU-seq (n = 2 independent experiments). (B) Bar charts showing the number of DE classes of transcripts in HeLa cells as assessed by 4sU-seq (n = 2 independent experiments). The degrees of differential expression are presented according to the legend. Conditions with (in hours) and without (-) 4-NQO or FP are shown. (C) Boxplots indicating the distribution of gene lengths for up-regulated and down-regulated protein-coding genes. Median gene length for each group is marked in grey and indicated above the boxes. (D) Top Molecular and Cellular Functions categories of the 4FP gene set as identified by IPA. The number of affected genes per category is shown on the right. (E) RT-qPCR of the indicated DNA damage-induced unspliced (pre-mRNA), uaRNA and eRNA transcripts. HeLa cells were treated as indicated by the legend. Results were normalized to the DMSO control and are presented as the mean ± s.e.m. (n = 3 independent experiments). *, P < 0.05; **, P < 0.01; ***, P < 0.001, determined by Student’s *t* test.

To explore further the role of P-TEFb in the DDR, we compiled a set of 4-NQO-induced coding genes, which showed decreased mRNA levels by at least 20 percent upon FP co-treatment (4FP set; Table S9). IPA identified cell death and survival, cellular development, cellular growth and proliferation, gene expression and cell cycle as the most significant cellular functions controlled by the 551-gene 4FP set (Figure 3D). Moreover, comparing it with gene sets of the Molecular Signatures Database (MSigDB) collection revealed that the 4FP set was enriched for nucleic acid binding proteins controlling gene expression (p-value=1.4×10^-16^-4.36×10^-33^; Table S10), which likely specify the response. In fact, the 4FP set bears similarity to the one controlled by NF-κB (p-value= 6.01×10^-57^) and of the p53 pathway (p-value=1.23×10^-40^) (Table S11). Notably, both transcription factors (TFs) function in DDR (Gomes et al., 2006; McKay et al., 2004; Sullivan et al., 2018; Wu et al., 2006) and require P-TEFb for inducing their target genes (Barboric et al., 2001; Gomes et al., 2006). The 4FP set overlaps significantly with the UV-induced gene sets of the MSigDB collection (p-value=3.8×10-50-2.02×10-56; Table S12) and is predicted by IPA to respond to DNA damage-inducing agents (Table S13).

We conducted kinetic quantitative RT-PCR (qRT-PCR) assays using HeLa cells to confirm that nascent transcripts of key DDR genes of the 4FP set, including oncogenic JUN, FOS and EGR1 (Lopez-Bergami et al., 2010; Thompson et al., 2009), a cell cycle inhibitor and a pro-survival CDKN1A/P21 (Besson et al., 2008; Cazzalini et al., 2010; Chen et al., 2009), an anti-apoptotic MCL1 (Zhou et al., 1997), and a DDR regulator GADD45β (Moskalev et al., 2012), are up-regulated in the wake of DNA damage-induced P-TEFb activation (Figure 3E). Importantly, co-administration of FP attenuated the induction of genes (Figure 3E). Similarly, the levels of oncogenic EGR2 pre-mRNA and its eRNA, as well as those of EXT1, RBM39 and their corresponding uaRNAs increased with a similar kinetic in a P-TEFb-dependent manner (Figure 3E). Together, these findings demonstrate that induced Pol II transcription of short protein-coding and non-coding genomic loci through P-TEFb is a hallmark of early DDR.

### RBM7 and 7SK snRNP enable induction of P-TEFb-dependent DDR genes

We next subjected the above protein-coding DDR genes to detailed mechanistic analyses. To provide further evidence that they are indeed controlled by P-TEFb upon genotoxic stress, we monitored the occupancy of CDK9 near transcription start sites (TSS) of the seven genes by conducting gene-specific quantitative chromatin immunoprecipitation (ChIP-qPCR) assays using unchallenged and 4-NQO-treated HeLa cells. Furthermore, we followed the occupancy of total Pol II and its P-TEFb-generated Ser2-P form near the TSS and in the middle of gene interior (INT) of each DDR gene. To take into account the expected 4-NQO-provoked changes in Pol II occupancy at the genes, we normalized Ser2-P signals to those of total Pol II. In accordance with our biochemical and nascent transcript findings, we observed that 4-NQO exposure elevated occupancy of CDK9 near the TSS of nearly all tested DDR genes (Figures 4A and S8A). Concomitantly, levels of the Ser2-P hallmark of productive Pol II elongation also increased at the sites upon genotoxic stress, particularly near TSS that correspond to the locations of paused Pol II (Figures 4A and S8A). Furthermore, FP blocked the increased Ser2-P levels (Figure S8B). The observed CDK9, Pol II and Ser2-P Pol II occupancies were specific as they were enriched significantly over the normal IgG and FOS intergenic site controls (Figure S8C). These results highlight further the role of P-TEFb in stimulating Pol II transcription following DNA damage.

**Figure 4.**
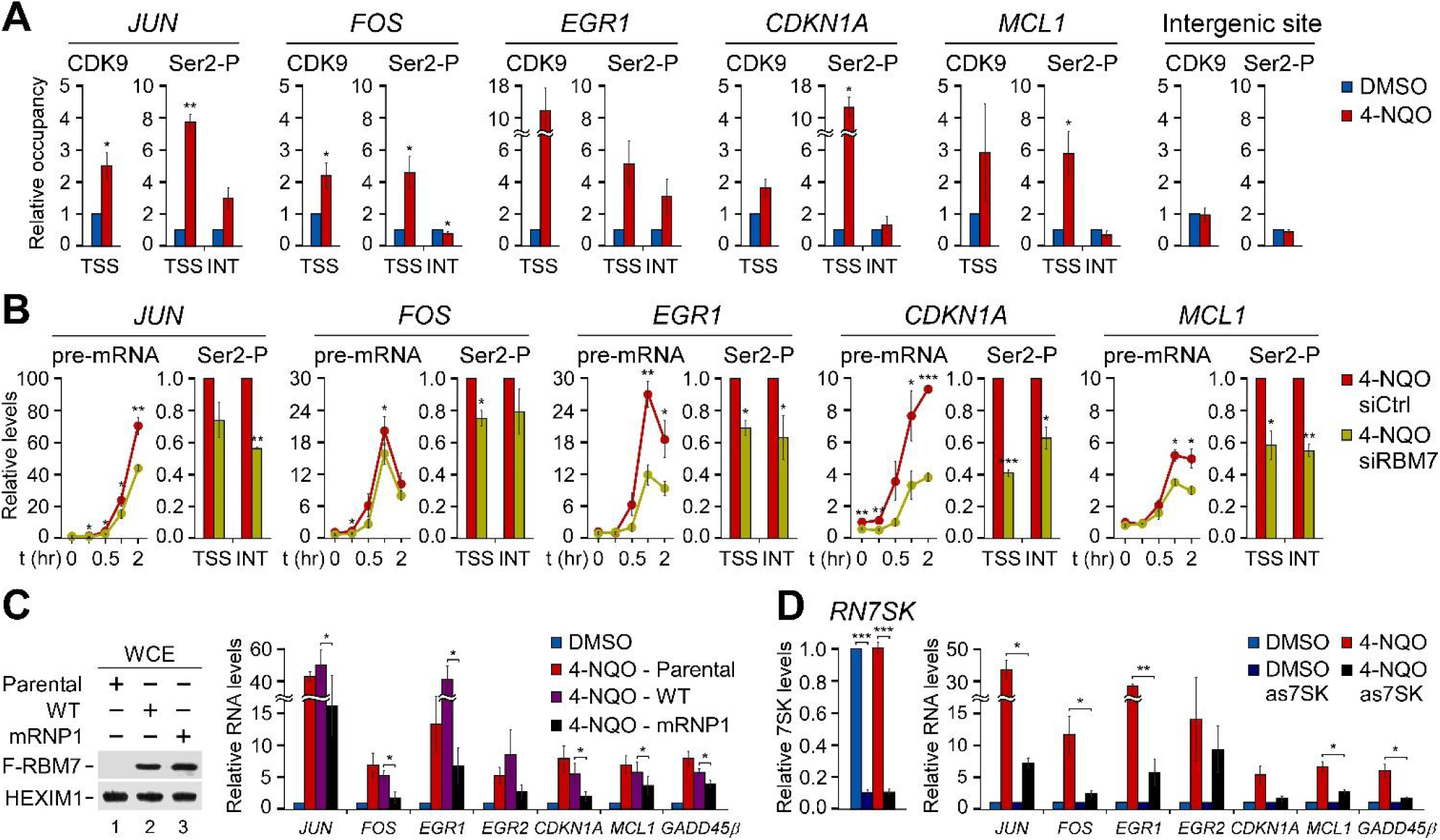
RBM7 and 7SK snRNP are critical for induction of P-TEFb-dependent DDR genes. (A) ChIP-qPCR of the occupancy of CDK9 and Ser2-P relative to Pol II at transcription start site (TSS) and in the middle of gene interior (INT) of the indicated DDR genes. The ChIP-qPCR data at the intergenic site approximately 100 kb upstream of *FOS* TSS is also presented. Conditions with (red bars) and without (blue bars) 4-NQO are shown. Results were normalized to the DMSO control and are presented as the mean ± s.e.m. (n = 3 independent experiments). *, P < 0.05; **, P < 0.01, determined by Student’s *t* test. (B) RT-qPCR (left) of unspliced transcripts (pre-mRNA) of the indicated DDR genes and ChIP-qPCR (right) of the levels of Ser2-P relative to Pol II at transcription start site (TSS) and in the middle of gene interior (INT) of the indicated DDR genes in control (siCtrl; red) and *RBM7* knockdown (siRBM7 #2; yellow) 4-NQO-treated HeLa cells. In RT-qPCR assays, the cells were exposed to 4-NQO for 15 min, 0.5 hr, 1 hr and 2 hr as indicated, and results were normalized to the untreated control and are presented as the mean ± s.e.m. (n = 3 independent experiments). *, P < 0.05; **, P < 0.01; ***, P < 0.001, determined by Student’s *t* test. ChIP-qPCR results were normalized to the control values that were set to 1 and are presented as the mean ± s.e.m. (n = 3 independent experiments). *, P < 0.05; **, P < 0.01, determined by Student’s *t* test. (C,D) RT-qPCR of unspliced transcripts of the indicated DDR genes in parental, WT and mRNP1 F-RBM7-expressing or 7SK-depleted HeLa cells. The cells were treated for 2 hr with DMSO or 4-NQO as indicated by the legend. Results were normalized to the respective DMSO control and are presented as the mean ± s.e.m. (C: n = 4 independent experiments; D: n = 3 independent experiments). *, P < 0.05; **, P < 0.01; ***, P < 0.001, determined by Student’s *t* test. Levels of the F-RBM7 proteins and efficacy of the *RN7SK* knockdown with the 7SK antisense DNA oligonucleotide (as7SK) are shown on the left.

Our findings thus far are consistent with the model in which following genotoxic stress, RBM7 activates P-TEFb by releasing it from the core of 7SK snRNP, resulting in transcriptional stimulation of specific gene sets. According to this scenario, interfering with RBM7 or its regulatory target, 7SK snRNP, should hamper the 4-NQO-triggered gene induction. Indeed, by using kinetic qRT-PCR assays we found that RBM7 knockdown led to the reduction of 4-NQO-induced levels of nearly all analyzed pro-survival and DDR gene transcripts in HeLa cells (Figures 4B and S8D). Importantly, this transcriptional defect coincided with decreased levels of the Ser2-P mark at the induced genes as revealed by ChIP-qPCR experiments (Figures 4B and S8D), underscoring the importance of RBM7 in enabling the P-TEFb-dependent gene induction. In further support of this notion, we found that HeLa cells constitutively expressing the 7SK binding-deficient mRNP1 F-RBM7 exhibited decreased induction of these genes upon 4-NQO exposure when compared to the parental or F-RBM7 expressing HeLa cells (Figure 4C), suggesting that mRNP1 F-RBM7 lowered the transcriptional response by acting in a dominant-negative fashion. Finally, we used 7SK-specific phosphorothioate-modified antisense DNA oligodeoxynucleotide to deplete the scaffolding 7SK, which disintegrates 7SK snRNP(Yik et al., 2003), and also detected a compromised transcriptional response to genotoxic stress (Figure 4D). Together, these results show the vital role of RBM7 in the DNA damage-induced activation of P-TEFb, which stimulates Pol II elongation at pro-survival and DDR genes.

### P-TEFb and RBM7 promote cell viability upon genotoxic stress

Given the importance of RBM7 and P-TEFb in facilitating transcriptional activation following genotoxic stress, we finally hypothesized that antagonizing the RBM7:P-TEFb axis during DDR should be detrimental to cell survival. This possibility was further supported by our analysis of the 4FP set with IPA Downstream Effects Analysis tool, which predicted that the genotoxic stress-induced genes under P-TEFb control promote cellular viability (Table S14). Thus, we performed time course cytotoxicity assays in which compromised membrane integrity of dead cells is proportional to fluorescence signal generated by the binding of an otherwise cell impermeable cyanine dye to DNA. For these experiments, we used HeLa cells, a cell line derived from the retinal pigment epithelium (RPE), and primary HFF-1 cells. Of note, UV-induced stress leads to death or dysfunction of RPE cells, which is thought to underlie several retinal diseases, including central vision loss of the age-related macular degeneration (Roduit and Schorderet, 2008). Confirming our hypothesis, we found that HeLa, RPE and HFF-1 cells became hypersensitive to 4-NQO when treated with the concentration of FP that attenuated the induction of P-TEFb-dependent target genes, reaching death of nearly all cells at 36 and 72 hours, respectively (Figures 5A and S9A). Depletion of RBM7 by two different siRNAs resulted in a similar hypersensitivity of HeLa and RPE cell lines to 4-NQO and UV (Figures 5B, S9B, and S9C). Of note, ectopically expressed F-RBM7, which was resistant to the siRNA-mediated repression targeting the 3’ untranslated region of the endogenous RBM7 transcripts, increased survival of 4-NQO-treated HeLa cells (Figure S9D). This provides further support that our RNAi findings were specific. We also conducted viability assays to provide additional evidence for the critical role of P-TEFb and RBM7 in the survival of genotoxic-stressed HeLa cells. We employed a fluorescence-based assay in which the indicator compound resazurin is reduced upon entering cells due to the reducing power of living cell cytosol, resulting in its fluorescence capacity. Corroborating our cytotoxicity results, P-TEFb inhibition and RBM7 depletion decreased the viability of 4-NQO-exposed HeLa cells (Figures S9E and S9F).

**Figure 5.**
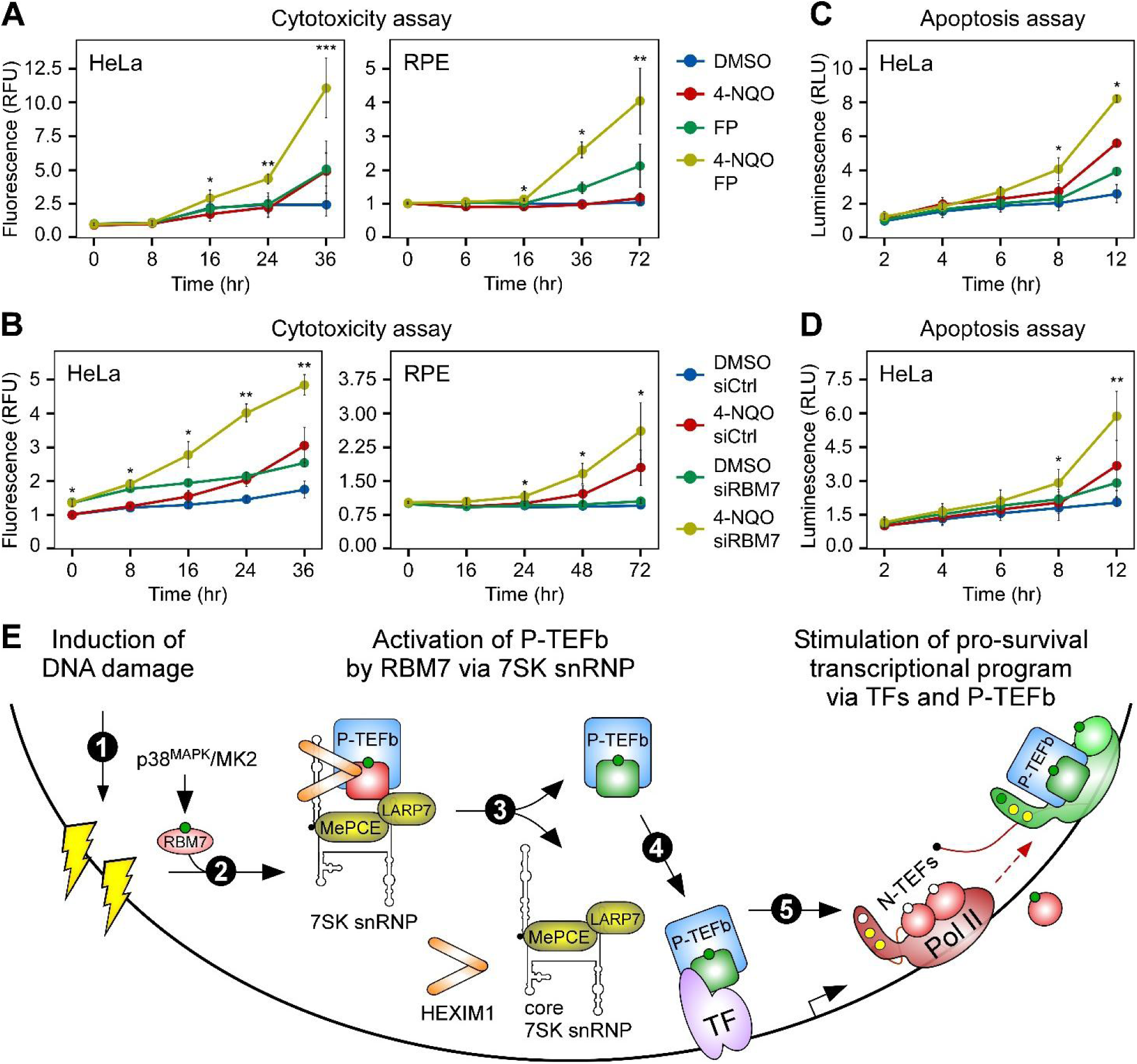
P-TEFb and RBM7 promote cell viability upon genotoxic stress. (A,B) Hypersensitivity of HeLa and RPE cells to 4-NQO upon FP treatment and RBM7 depletion. The cells were treated as indicated by the legends and examined at the time points indicated below the graphs. Two independent siRNAs (siRBM7 #2, HeLa cells; siRBM7 #1, RPE cells) were used to deplete RBM7. Cytotoxicity results are presented as fluorescence values relative to the untreated control and plotted as the mean ± s.e.m. (n = 3 independent experiments). *, P < 0.05; **, P < 0.01; ***, P < 0.001, determined by Student’s t test using 4-NQO and 4-NQO FP or 4-NQO siRBM7 data sets, respectively. (C,D) FP treatment and RBM7 depletion enhance 4-NQO-induced apoptosis in HeLa cells. The cells were treated as indicated by the legends and examined at the time points indicated below the graphs. siRBM7 #2 was used to deplete RBM7. Apoptosis results are presented as luminescence values relative to the untreated control and plotted as the mean ± s.e.m. (n = 3 independent experiments). *, P < 0.05; **, P < 0.01, determined by Student’s t test using 4-NQO and 4-NQO FP or 4-NQO siRBM7 data sets, respectively. (E) Model of P-TEFb activation by RBM7 during DDR. DNA damage signaling (step 1) provokes phosphorylation (green circle) of RBM7 by the p38MAPK/MK2 pathway. Subsequently, RBM7 of the trimeric NEXT complex binds 7SK snRNP (step 2) to release the inactive P-TEFb (CDK9 in red) from the core of 7SK snRNP (step 3), yielding active P-TEFb (CDK9 in green). In turn, transcription factors (TF) capture P-TEFb on chromatin (step 4). Stimulation of pro-survival DDR gene transcription at the Pol II pause release phase ensues (step 5), which is achieved by P-TEFb-mediated phosphorylation (green circles) of Pol II CTD at Ser2 as well as the negative transcription elongation factors (N-TEFs) NELF and DSIF. While NELF dissociates from Pol II, DSIF becomes a positive transcription elongation factor.

Finally, we asked if apoptosis was responsible for the hypersensitivity of P-TEFb or RBM7 deficient cells to genotoxic stress. Thus, we subjected HeLa cells to the same conditions as in the above functional experiments. We followed kinetics of apoptosis using a luminescence-based assay that measures the exposure of phosphatidylserine on the outer cell membrane surface during apoptosis with two Annexin V fusion proteins containing fragments of NanoBit Luciferase (Dixon et al., 2016), of which complementation is achieved through Annexin V - phosphatidylserine binding. Indeed, FP treatment and RBM7 depletion enhanced 4-NQO-stimulated apoptosis (Figures 5C and 5D). Comparison of the cytotoxicity and apoptosis kinetics shows that the onset of apoptosis precedes the occurrence of cell death, indicating that apoptosis was causing the cellular demise. As an independent measure of apoptosis activation, we also monitored cleavage of poly(ADP-ribose) polymerase (PARP) and Caspase-3 by Western blotting. Supporting our kinetic results, we also found that inhibiting P-TEFb activation in 4-NQO-treated HeLa cells with FP and RBM7 depletion enhanced the levels of PARP and Caspase-3 cleavage products (Figures S9G and S9H). Together, these findings show the critical importance of the RBM7:P-TEFb axis in enabling a pro-survival transcriptional response to genotoxic stress.

## DISCUSSION

Reshaping cellular gene expression program in the face of external or internal stress is a fundamental characteristic of all living forms. In this study, we investigated the mechanisms and importance of transcriptional activation following genotoxic stress. We uncovered that stimulation of Pol II pause release by P-TEFb is at the heart of cellular DDR (Figure 5E). We identified RBM7 of the trimeric NEXT complex as a critical regulator directing the DNA damage-induced release of P-TEFb from the core of 7SK snRNP, which in turn enables a concerted transcriptional response that is indispensable to the survival of stressed cells. We envision that upon activation of P-TEFb, its recruiting factors including Brd4, the Mediator and transcriptional activators (Quaresma et al., 2016; Takahashi et al., 2011; Yang et al., 2005), especially those playing critical roles in DDR such as NF-κB and p53 (Barboric et al., 2001; Gomes et al., 2006; McKay et al., 2004; Wu et al., 2006), capture the kinase on chromatin, thereby underwriting specificity to the transcriptional outcome and resolution to the genotoxic affront.

Our work provides a novel paradigm in stress-dependent control of Pol II pause release. The exosome-independent role of RBM7 in the activation of P-TEFb expands our knowledge of its regulatory ability, which has been thus far limited to enabling RNA degradation (Lubas et al., 2015). Considering that UV irradiation impairs the RBM7-mediated ribonucleolysis by triggering its phosphorylation via the p38MAPK/MK2 pathway (Blasius et al., 2014; Tiedje et al., 2015), genotoxic stress thus switches a function of RBM7 from facilitating RNA degradation to promoting DNA transcription. Of note, RBM7 binds near-stoichiometric levels of hMTR4 and ZCCHC8 in cell extracts (Andersen et al., 2013; Lubas et al., 2011), indicating its primary engagement within NEXT. Therefore, despite the capacity of purified RBM7 to release P-TEFb from 7SK snRNP in vitro, it is likely that RBM7 accomplishes this task in vivo as a subunit of NEXT. This possibility could explain the observed dominant-negative effect of the 7SK binding-deficient mRNP1 F-RBM7 on the induction of DDR genes, where the mutant protein might interfere with the function of endogenous RBM7 by dislodging it from NEXT. We speculate that P-TEFb activation is not limited to DDR. It might take place also in response to other types of stress, such as hypoxic milieu of tumor cells, in which the oncogenic transcriptional activator HIF1A could co-opt increased availability of active P-TEFb for alleviating Pol II pausing at hypoxia-inducible genes (Galbraith et al., 2013).

Importantly, this study clarifies our understanding of the significance of stimulating Pol II transcription upon DNA damage. Since P-TEFb targets promoter-proximal Pol II that is poised for productive elongation, its stress-induced activation and chromatin recruitment is suited uniquely to promote the rapid and coordinated transcriptional response. Our 4sU-seq and ChIP-qPCR findings are in agreement with recent studies reporting increased levels of nascent RNA synthesis at promoter-proximal regions following UV irradiation (Andrade-Lima et al., 2015; Lavigne et al., 2017; Williamson et al., 2017). It remains to be established, however, how exactly does the stimulation of Pol II pause release via RBM7 and P-TEFb promote survival of cells experiencing genotoxic stress. We propose that transcriptional reprogramming plays a pivotal role. While induction of coding DDR genes as exemplified by CDKN1A/P21 and MCL1, as well as lincRNAs enables cell cycle arrest and opposes apoptosis (Cazzalini et al., 2010; Huarte et al., 2010; Hung et al., 2011; Zhou et al., 1997), elevated levels of eRNAs and uaRNAs might concurrently stabilize new gene expression program through various mechanisms, which could include local modulation of chromatin environment (Bose et al., 2017; Lai et al., 2013; Mousavi et al., 2013), facilitation of TFs occupancy (Sigova et al., 2015), and augmentation of Pol II pause release (Schaukowitch et al., 2014). Accordingly, we speculate that the transiently reshaped transcriptome sets the stage for subsequent recovery from genotoxic stress. Therein activation of P-TEFb could be also advantageous to the repair of damaged DNA itself. Supporting this premise, stimulating the release of Pol II from promoter-proximal pausing could enhance detection of DNA damage throughout the transcribed genome, resulting in more effective transcription-coupled NER (TC-NER) (Andrade-Lima et al., 2015; Chiou et al., 2017; Lavigne et al., 2017). Namely, Pol II stalled at the sites of DNA damage mediates activation of this pathway (Hanawalt and Spivak, 2008; Vermeulen and Fousteri, 2013), and because Pol II dissociates from DNA during individual TC-NER reactions (Chiou et al., 2017), the repair of following DNA lesions would benefit from successive rounds of Pol II pause release events. In addition, active P-TEFb might exert its positive role via its DNA damage-induced interaction with Cockayne syndrome B translocase (Boeing et al., 2016), which is instrumental for TC-NER and resetting of Pol II transcription as cells recover from DNA damage (Epanchintsev et al., 2017; Mayne and Lehmann, 1982).

Finally, P-TEFb-dependent Pol II pause release is frequently dysregulated in cancers, particularly in those addicted to c-MYC and translocations of mixed-lineage leukemia gene (Dawson et al., 2011; Delmore et al., 2011; Huang et al., 2014; Ji et al., 2014; Luo et al., 2012; Rahl et al., 2010; Tan et al.), spurring interest in the development of highly specific CDK9 inhibitors for clinical use (Lu et al., 2015; Olson et al., 2018). Our discovery of the pro-survival role of P-TEFb in genotoxic stress reinforces the relevance of this critical transcriptional kinase to cancer biology. Hence, our work offers a framework for novel anti-cancer approaches with targeting RBM7 and P-TEFb in addition to DNA-damaging chemotherapeutics. Countering the RBM7:P-TEFb axis might prove especially effectual in combination with broad spectrum bulky DNA adduct-inducing platinum-based drugs such as cisplatin, which shows improved therapeutic outcome in NER-deficient cancers (Dietlein et al., 2014; Gavande et al., 2016). The proposed combinatorial approach might also curb the emergence of drug-resistant tumor cells (Sharma et al., 2010; Shen et al., 2012; Yashiro-Ohtani et al., 2014), a major challenge of contemporary cancer monotherapies.

## EXPERIMENTAL PROCEDURES

### Cell Culture

HEK 293, HEK 293 Flp-In T-REx (Thermo Fisher Scientific), HeLa (ATCC), HeLa Flp-In and human foreskin fibroblast (HFF-1, ATCC) cell lines were grown in Dulbecco‘s Modified Eagle‘s Medium (D-MEM; Sigma, D5796) supplemented with 10% fetal bovine serum (FBS) and 100 U/ml penicillin/streptomycin. The parental HEK 293 Flp-In T-REx and HeLa Flp-In cell lines were grown with 50μg/ml of zeocin (InVivogen, ant-zn-1). HEK 293 Flp-In T-REx cell lines expressing 3X-FLAG peptide or 3X-FLAG epitope-tagged proteins were generated according to the manufacturer’s instructions and grown with 100 μg/ml of Hygromycin B Gold (InVivogen, ant-hg-5) and 3 μg/ml of Blasticidin (InVivogen, ant-bl-1). HeLa Flp-In F-CBP20 cell line was grown with 200 μg/ml of Hygromycin B Gold. Both HeLa Flp-In cell lines were a gift from Dr. Bertrand and described previously (Hallais et al., 2013). Retinal pigment epithelium (RPE) cell line (ATCC) was grown in D-MEM/F12 (Sigma, D8437) supplemented with 10 % fetal bovine serum (FBS) and 100 U/ml penicillin/streptomycin. Cell lines were maintained at 37 °C with 5 % CO_2_. Cell lines used in this study are listed in Supplementary Table 15.

### Plasmid DNAs and Mutagenesis

To generate plasmid DNAs encoding 3X-FLAG epitope-tagged HEXIM1, LARP7, CDK9 and RBM7, the cDNAs were amplified using Phusion DNA polymerase (NEB, M0530) with primers carrying the appropriate restriction enzymes sites and cloned using Rapid Ligation Kit (Thermo Fisher Scientific, K1423) into pcDNA5/FRT/TO (https://www.addgene.org/vector-database/2133/) vector, which was modified to encode 3X-FLAG peptide upstream of the multiple cloning site (a gift from Dr. Ule). To generate plasmids encoding the mutant RBM7 proteins, pcDNA5/FRT/TO/3X-FLAG-RBM7 was used for mutagenesis with Q5^®^ Site-Directed Mutagenesis Kit (NEB, E0554S). To generate pcDNA5/FRT/TO/3X-FLAG-RBM7 mRNP1, the primer sequences used were: 5’-TGCGGCTGTGAATTTCAAACATGAAGTG-3’; 5’-GCCTGCGCTGGTTTACCATCCTTATCTTTTG-3’. To generate pcDNA5/FRT/TO/3X-FLAG-RBM7 S136A, the primer sequences used were: 5’-GCTTCTCCAGAAAATTTTCAGAGACAAG-3’; 5’-GAAAGATCTCTGAATTATCTGTGCTG-3’. Plasmids used in this study are listed in Supplementary Table 16.

### Chemicals and Treatments

4-Nitroquinoline N-oxide (4-NQO; Sigma, N8141) was diluted in DMSO to a final concentration of 50mM, aliquoted, sealed and stored at −80 °C. UV irradiation was performed in Crosslinker CL-1000 using 254 nm wavelength lamp with dose 40-60 J/m^2^. Flavopiridol (Sigma, F3055) and 4sU (Sigma, T4509) were diluted in DMSO to a final concentration of 1 mM and stored at −20 °C. Tetracycline hydrochloride (Sigma, T7660) was diluted in water to a final concentration of 1mg/ml, aliquoted and stored at −20 °C.

### RBM7 iCLIP Assay

HEK 293 Flp-In T-REx F-RBM7 cells were treated for 24 hours with 2 μg/ml of doxycycline to express F-RBM7, exposed for 2 hours to DMSO or 5 mM of 4-NQO, washed once with PBS and UV-cross-linked at 0,15 mJ/cm^2^ with 254 nm wavelength. Retrieval of protein-bound RNAs and preparation of cDNA libraries were prepared as described previously (Huppertz et al., 2014). Cross-linked nucleotides were defined as the nucleotide upstream of mapped iCLIP cDNA tags as described previously (Konig et al., 2010). Replicate iCLIP experiments were performed, cross-linking positions compared between samples, and the replicates were subsequently combined into groups for final analysis using the iCount package (http://icount.biolab.si/).

### Quantitative RNA Immunoprecipitation Assay

HEK 293 Flp-In T-REx cells grown on 15 cm plates were treated for 16 hours with 1 μg/ml of tetracycline to express the 3X-FLAG epitope-tagged proteins and exposed at approximately 90 % confluency to DMSO or 10 μM of 4-NQO for 2 hours. The cells were resuspended in Falcon tubes with 10 ml of ice-cold PBS and cross-linked with 1% formaldehyde at room temperature for 10 minutes, which was stopped with 250 mM of glycine for 5 minutes. The cells were then washed twice with ice-cold PBS, resuspended in buffer A (5 mM PIPES, 85 mM KCL, 0.5 % NP-40, pH 7.9), and incubated 10 minutes on ice for nuclear extraction. Nuclear pellets obtained by centrifugation were lysed in 1 ml of RIPA buffer (50 mM Tris pH 8.0, 150 mM NaCL, 5 mM EDTA pH 8.0, 0.5 % sodium deoxycholate, 1 % NP-40, 0.1 % SDS) in the presence of EDTA-free Protease Inhibitor Cocktail (Sigma, 11873580001) and SUPERase In RNase Inhibitor (Thermo Fisher Scientific, AM2694). The lysates were then sonicated during one round of 35 cycles of 30 sec ON/30 sec OFF at 4 °C with the Bioruptor Plus sonication device (Diagenode, B01020001) combined with the Bioruptor Water cooler (Cat. No. BioAcc-cool) & Single Cycle Valve (Cat. No. VB-100-0001) at high power setting (position H) using 1.5 ml TPX microtubes (Cat. No. M-50001). After centrifugation at 13 000 g for 15 min, 5 % of supernatant was stored at - 80 °C for determining RNA input. The rest of the sample was divided in three equal parts, which were supplemented with additional 600 μl of RIPA buffer and incubated overnight at 4 °C with 15 μl of antibody-coupled protein G Dynabeads (Thermo Fisher Scientific, 10004D). Before adding the supernatant, the beads were pre-blocked with bovine serum albumin and salmon sperm DNA overnight at a final concentration of 0.2 μg/μl, pre-incubated in 500 μl of RIPA buffer for 4 hours with the antibody and collected by magnetic stand to remove the unbound antibody. We used 3 μg of normal mouse IgG (Santa Cruz Biotechnology, sc-2025), 3 μg of anti-FLAG (Sigma, F1804), or 3 μg of anti-HEXIM1 (Everest Biotech, EB06964) antibody. The beads were then washed with RIPA high salt buffer (20 mM Tris pH 8.0, 500 mM NaCL, 2 mM EDTA pH 8.0, 1 % Triton-X, 0.1 % SDS), RIPA low salt buffer (20 mM Tris pH 8.0, 150 mM NaCL, 2 mM EDTA pH 8.0, 1 % Triton-X, 0.1 % SDS), LiCl wash buffer (250 mM LiCl, 1 % NP-40, 1 % sodium deoxycholate) and TE buffer (10 mM Tris, 1 mM EDTA, pH 8.0). RNA-protein complexes were eluted at room temperature with elution buffer (1 % SDS, 100 mM NaHCO_3_). After reverse the cross-linking, RNAs from eluates and inputs were isolated using TRIzol LS reagent (Thermo Fisher Scientific, 10296028) according to the manufacturer’s protocol. RNA samples were DNase-treated with the Turbo DNA-Free kit (Ambion, AM1907), reverse transcribed with SuperScript III reverse transcriptase (Thermo Fisher Sceintific, 18080044) and random hexamers (Thermo Fisher Scientific, N8080127), and amplified using FastStart Universal SYBR Green QPCR Master (Rox) (Roche, 04913914001), RNA-specific primer pair, and Stratagene Mx3005 qPCR machine. Primers were from Integrated DNA Technologies and designed using PrimerQuest Tool. Values were normalized to their levels in RNA inputs and calculated using the MxPro QPCR Software. Results from at least three independent experiments are presented as the mean ± s.e.m. Sequences of the primers used in RIP-qPCR assays are listed in Supplementary Table 17.

### Co-immunoprecipitation and Western Blotting Assays

HEK 293 Flp-In T-REx cells grown on 10 cm plates were treated for 16 hours with 1 μg/ml of tetracycline to express the 3X-FLAG epitope-tagged proteins and exposed to DMSO or 10 μM of 4-NQO for 2 hours. Whole cell extracts (WCE) were prepared by lysing the cell pellets on ice for 15 min with buffer C (20 mM Tris-HCl, 0.5 % NP-40, 150 mM NaCl, 1.5 mM MgCl_2_, 10 mM KCl, 10 % Glycerol, 0.5 mM EDTA, pH 7.9) in the presence of protease cocktail inhibitor (Thermo Fisher Scientific, 88266), followed by 10 seconds sonication with the Misonix XL-2000 Ultrasonic Liquid Processor using the P-1 Microprobe 3.2 mm tip, power setting 3, and centrifugation of lysates at 20 000 g for 15 min. For FLAG immunoprecipitation, WCE were incubated at 4 °C for 4 hours with buffer C-equilibrated anti-FLAG M2 affinity gel (Sigma, A2220), immuno-complexes were washed three times with buffer C, eluted in SDS running buffer in the presence of dithiothreitol (DTT-RO, Sigma) for 5 min at 95 °C and resolved using 12% SDS-PAGE. For total Pol II co-immunoprecipitation (co-IP), 1 μg of the anti-Pol II 8WG16 antibody (Abcam Biotech, ab817) was immobilized on Protein G Dynabeads (Thermo Fisher Scientific, 10004D) according to manufacturer’s instructions. WCE were prepared using ELB buffer (50 mM HEPES-KOH pH 7.9, 0.1% Triton X-100, 5 mM dithiothreitol, 5 mM EDTA, 150 mM NaCl), incubated with the ELB buffer-equilibrated beads for 16 hours at 4 °C, and the beads were washed three times ELB buffer. For endogenous HEXIM1 co-IP, 2 μg of the anti-HEXIM1 antibody (Everest Biotech, EB06964) was immobilized on Protein G Dynabeads (Thermo Fisher Scientific, 10004D) according to manufacturer’s instructions. In the case of HEXIM1 co-IP using HEK 293 Flp-In T-REx F-RBM7 and HEK 293 Flp-In T-REx F-RBM7 mRNP1 cell lines, the cells were treated for 24 hours with 1 μg/ml of tetracycline to express the 3X-FLAG epitope-tagged proteins. Upon cell lysis in buffer C, WCE were then added to the buffer C-equilibrated beads and incubated for 16 hours at 4 °C, and the beads were washed three times with buffer C. 50 % of the beads was used to determine the levels of HEXIM1, CDK9 and F-RBM7 by Western blotting. To determine the levels of 7SK in anti-HEXIM1 co-IP, 500 μl of TRI reagent (Sigma, T9424) was added to the remaining 50 % of the beads for RNA extraction. RNA samples were DNase-treated with the Turbo DNA-Free kit (Ambion, AM1907), reverse transcribed with SuperScript III reverse transcriptase (Thermo Fisher Sceintific, 18080044) and random hexamers (Thermo Fisher Scientific, N8080127), and quantified with the Stratagene Mx3005 qPCR machine as described above using 5S rRNA as a normalizing control. Results from independent experiments are presented as the mean ± s.e.m. For Western blotting, the following antibodies were used according to manufacturers’ instructions: anti-FLAG (Sigma, F3165); anti-RBM7 (Sigma, HPA013993; Proteintech, 21896-1-AP); anti-HEXIM1 (Everest Biotech, EB06964); anti-CDK9 (Santa Cruz Biotechnology, sc-484); anti-LARP7 (a gift from Dr. Qiang Zhou described previously(He et al., 2008)); anti-MePCE (Santa Cruz Biotechnology, sc-82542); anti-RNA polymerase II CTD repeat YSPTSPS (phospho S2) antibody (Abcam, ab5095); anti-hnRNP A1 (Abnova, MAB2492); anti-*γ*-H2AX (Abcam, ab22551); anti-hMTR4/SKIV2L2 (Abcam, ab70551); anti-EXOSC3 (Santa Cruz Biotechnology, sc-166568); anti-cyclin T1 (Santa Cruz Biotechnology, sc-10750), anti-Cleaved PARP (Cell Signaling Technology, 9541); anti-Cleaved Caspase-3 (Cell Signaling Technology, 9661). Manufacturers provide validation for all antibodies.

### Immunofluorescence Microscopy

HeLa cells were grown on polylysine coated coverslips, treated with DMSO or 10 μM of 4-NQO for 1 hour, washed twice in PBS, and incubated in cytoskeletal buffer (CSK; 10 mM Pipes pH 6.8, 300 mM sucrose, 100 mM NaCl, 3 mM MgCl_2_, 1 mM EGTA) for 10 min. Cells were then fixed in 4% formaldehyde in CSK buffer for 1 hour on ice and blocked with TBS-I (10 mM Tris pH 7.7, 150 mM NaCl, 3 mM KCl, 1.5 mM MgCl_2_, 0.05% Tween 20, 0.1% BSA, 0.2% glycine) for at least 1 hour on ice. Primary antibody staining using anti-*γ*H2AX (Abcam, ab22551; 1:100 dilution) was done overnight at 4 °C, which was followed by two washes with ice cold CSK buffer. Secondary antibody staining using goat anti-mouse conjugated with AlexaFluor 488 (Thermo Fisher Scientific, A11001; 1:1000 dilution) was performed for 4 h at 4 °C. Cells were then washed twice with ice cold CSK buffer, incubated with NucBlue reagent (R37606, Thermo Fisher Scientific) for 30 min at room temperature and washed three times with ice cold CSK buffer. Coverslips were mounted with ProLong Gold Antifade Mountant (Thermo Fisher Scientific, P10144). Images were acquired by AxioLab microscope equipped with AxioCam and AxioVision 4.3 software (Zeiss), and analyzed using CorelDRAW Graphic Suite 2017. Representative images show cells treated with DMSO or 250 nM of 4-NQO for 1 hour. The number of cells with at least one nuclear γ-H2AX foci from two independent experiments was counted, plotted as percentage of the total number of cells in the field, and presented as the mean ± s.e.m. For each independent treatment, cells were counted from at least ten fields containing at least ten cells per field.

### Glycerol Gradient Sedimentation Analysis

HeLa Flp-In cells were treated with DMSO or 10 μM of 4-NQO for 2 hours, lysed on ice for 15 min in 0.6 ml of lysis buffer B (20 mM HEPES, 0.3 M KCl, 0.2 mM EDTA, 0.1 % NP-40, 0.1 % protease inhibitor, pH 7.9). Cellular extracts were then subjected to ultracentrifugation in a SW41 Ti rotor (Beckman) at 38 000 rpm for 16 hours in a 10 ml glycerol gradient solution (10–30 %) containing buffer B. Fractions were collected and analyzed as described previously (Yik et al., 2005).

### Bimolecular Fluorescence Complementation Assay

HeLa Flp-In cells grown on 6-well plates were co-transfected with 0.2 μg of pEF.YN-CTD and 2 μg of pEF.YC-P-TEFb plasmids using X-tremeGENE transfection reagent (Sigma, XTG9-RO). Twenty-four hours after transfection, the cells were seeded into 6-8 wells of a 24-well plate and grown an additional 24 to 48 hours. The cells were then left untreated or treated with DMSO, with 2, 5, and 10 μM of 4-NQO for 0.5, 1, and 4 hours, and with 5 μM of SAHA or JQ1 for 1 hour. Fluorescence signals were detected using Olympus IX70 fluorescent microscope. The fluorescence images were analyzed using MetaMorph software. YFP positive cells were counted manually and averaged from three randomly chosen fields of each sample.

### P-TEFb Release Assay

P-TEFb release assay was performed as described previously (Calo et al., 2015) with the following modifications. WCE from one confluent 15 cm plate of HeLa cells was prepared by lysing the cell pellets on ice for 30 min with 1.2 ml of buffer C in the presence of protease cocktail inhibitor (Thermo Fisher Scientific, 88266). To immobilize 7SK snRNP, 1 ml of WCE was incubated at 4 °C for 4 hours with 75 μl of Protein G Dynabeads (Thermo Fisher Scientific, 10004D) that were pre-bound with 5 μg of anti-HEXIM1 antibody (Everest Biotech, EB06864). For the control, the remaining 200 μl of WCE was incubated with 15 μl of Protein G Dynabeads (Thermo Fisher Scientific, 10004D) that were pre-bound with 1 μg of normal IgG antibody. Immuno-purified F-RBM7 proteins were incubated with the equal amounts of immobilized 7SK snRNP for 2 hours on ice, which was followed by the collection of samples for Western blotting analysis from 50 % of the beads. To determine the levels of 7SK that remained bound to HEXIM1, 500 μl of TRI reagent (Sigma, T9424) was added to the remaining 50 % of the beads for RNA extraction. RNA samples were DNase-treated with the Turbo DNA-Free kit (Ambion, AM1907), reverse transcribed with SuperScript III reverse transcriptase (Thermo Fisher Sceintific, 18080044) and random hexamers (Thermo Fisher Scientific, N8080127), and quantified with the Stratagene Mx3005 qPCR machine as described above using 5S rRNA as a normalizing control. Results from independent experiments are presented as the mean ± s.e.m. For purification of the 3X-FLAG epitope-tagged RBM7 proteins, the corresponding HEK 293 Flp-In T-REx F-RBM7 cells grown on three 15 cm plates were treated for 16 hours with 1 μg/ml of tetracycline to express the F-RBM7 proteins. The cells were lysed in buffer C. WCE were then incubated with 30 μl of buffer C-equilibrated anti-FLAG M2 affinity gel (Sigma, A2220) for 16 hours in the presence of protease inhibitor (Thermo Fisher Scientific, 88665) and RNase A (Thermo Fisher Scientific, 12091021). After incubation, samples were washed three times with buffer C, where the first two washes contained 600 mM of NaCl. F-RBM7 proteins were eluted with 300 μg/ml of 3X-FLAG peptide (ApexBio Technology, A6001) in 50 μl of buffer C, and 10 μl (+) or 20 μl (++) of the eluates were used per each reaction. To examine their purity, 10 μl eluate samples were subjected to SDS-PAGE and Western blotting using anti-hMTR4, anti-EXOSC3, and anti-CDK9 antibodies.

### 4sU-sequencing Assay

Metabolic labeling and isolation of newly transcribed RNA was performed as described previously (Radle et al., 2013). In brief, 80 % confluent Hela Flp-In cells were mock-treated or treated with DMSO, 5 μM of 4-NQO, or 5 μM of 4-NQO and 250 nM of flavopiridol for 1 or 2 hours before lysis. Thirty minutes prior to lysis, the cells were labeled with 4sU at a final cell culture medium concentration of 100μM. RNA was extracted with TRIzol reagent (Thermo Fisher Scientific, 15596018) and 150 μg of total RNA was used for biotinylation with EZ-Link Biotin-HPDP (Thermo Fisher Scientific, 21341) for 90 minutes at room temperature. Second round of RNA extraction was performed with chloroform-isopropanol. µMACS Streptavidin Kit (Miltenyi, #130-074-101) was used for separation of labeled RNA, which was followed by elution with DTT and RNA extraction with isopropanol. Libraries from two biological replicates were prepared and sequenced by Beijing Genomics.

### Analysis of 4sU-sequencing Data Sets

Sequencing quality was assessed with FastQC (http://www.bioinformatics.babraham.ac.uk/projects/fastqc/) and sequencing adapters were trimmed using Cutadapt (Martin, 2011). Reads were mapped against the human reference genome (GRCh38/hg38) and rRNA sequences using ContextMap 2 (Bonfert et al., 2015). Read counts for mRNAs, lincRNAs, uaRNAs and eRNAs were calculated using featureCounts (Liao et al., 2014). mRNA and lincRNA annotation were taken from Gencode version 25 and eRNA annotations were taken from the study by Lubas et al. (Lubas et al., 2015). eRNAs within 5 kb of an annotated gene (according to Gencode) were excluded. uaRNAs were defined as the window from −3kb to the transcription start sites of mRNAs and lincRNAs on the opposite DNA strand. If according to the Gencode annotation another gene was present within 10 kb upstream of the uaRNA-associated gene, we excluded those uaRNAs from our analysis. Expression of mRNAs, lincRNAs, uaRNAs and eRNAs was quantified in terms of fragments per kilobase of exons per million mapped reads (FPKM) and averaged between replicates. Differential gene expression analysis to determine fold-changes in gene expression and significance of changes was performed using edgeR (Robinson et al., 2010). Here, only RNAs with an average read count ≥1 in the respective samples were included. EdgeR models read counts using the negative binomial distributions and uses the quantile-adjusted conditional maximum likelihood method to estimate log fold-changes and p-values for differential gene expression. P-values obtained from edgeR were corrected by multiple testing using the method by Benjamini and Hochberg (Benjamini and Hochberg, 1995) for adjusting the false discovery rate (FDR) and a p-value cutoff of 0.01 was applied. For correlation analysis, Spearman rank correlation was used. Gene lengths were obtained from Gencode annotations. Supplementary tables 3–6 contain mRNAs, lincRNAs, uaRNAs and eRNAs which were differentially expressed upon 1 and 2 hours of 4-NQO treatment. A transcript was considered consistently regulated if (i) the FDR-adjusted p-value was ≤ 0.01 in at least one of the two experimental conditions, (ii) it was regulated in the same way (either up or down) in both experimental conditions and (iii) it was not differentially expressed in the DMSO control condition.

### Ingenuity Pathway Analysis and Molecular Signature Database Analysis

The set of protein-coding genes that are differentially expressed upon 1 and 2 hours of 4-NQO exposure and the 4FP gene set were subjected to IPA Upstream Regulator Analysis, which identifies upstream regulators that can explain the observed gene expression changes in a user’s dataset. The 4FP gene set was also subjected to IPA Downstream Effects analytic, which identifies biological functions that are expected to be increased or decreased given the observed gene expression changes in a user’s dataset. Detailed explanation of these analyses is provided by IPA at http://pages.ingenuity.com/rs/ingenuity/images/0812%20upstream_regulator_analysis_whitepaper.pdf; and http://pages.ingenuity.com/rs/ingenuity/images/0812%20downstream_effects_analysis_whitepaper.pdf. Overlaps between the 4FP gene set and gene sets of the Molecular Signatures Database (MSigDB) collection v6.0 were computed using the on-line tool at http://software.broadinstitute.org/gsea/msigdb/ index.jsp. Gene set enrichment analysis, GSEA software, and MSigDB were described previously (Subramanian et al., 2005).

### RNA extraction and RT-qPCR Analysis

Eighty % confluent Hela Flp-In cells grown on 6-well plates were left untreated or treated with DMSO, 5 μM of 4-NQO, or 5 μM of 4-NQO and 250 nM of flavopiridol for 15 min, 30 min, 1 hour and 2 hours. RNA samples were extracted using TRI Reagent (Sigma, T9424), DNase-treated with the Turbo DNA-Free kit (Ambion, AM1907), and reverse transcribed with M-MLV reverse transcriptase (Thermo Fisher Scientific, 28025-013) and random hexamers (Thermo Fisher Scientific, N8080127) according to the manufacturers’ instructions. qPCR reactions were performed with diluted cDNAs, primer pairs that spanned exon-intron junction, and FastStart Universal SYBR Green QPCR Master (Rox) (Roche, 04913914001) using Stratagene Mx3005 qPCR machine. Primers were from Integrated DNA Technologies and designed using PrimerQuest Tool. Relative levels of transcripts were calculated using the MxPro QPCR Software v4.10. For 7SK, 7SK DNA value was used as a normalizer. For other RNAs, *GAPDH* mRNA values were used as a normalizer. Results from at least three independent experiments are presented as the mean ± s.e.m. Sequences of the primers used in RT-qPCR assays are listed in Supplementary Table 18.

### Quantitative Chromatin Immunoprecipitation Assay

ChIP-qPCR assay was performed as described previously (Ekumi et al., 2015) with the following modifications. HeLa Flp-In cells grown on 15 cm plates were treated at approximately 90 % confluency with DMSO or 5 μM of 4-NQO for 2 hours. For the total Pol II and Ser2-P Pol II assays, the formaldehyde cross-linked cell pellets were lysed in 800 μl of RIPA buffer (50 mM Tris pH 8.0, 150 mM NaCL, 5 mM EDTA pH 8.0, 0.5 % sodium deoxycholate, 1 % NP-40, 0.1 % SDS) in the presence of EDTA-free Protease Inhibitor Cocktail (Sigma, 11873580001) and SUPERase In RNase Inhibitor (Thermo Fisher Scientific, AM2694). Lysates were then sonicated during one round of 35 cycles of 30 sec ON/30 sec OFF at 4 °C with the Bioruptor Plus sonication device (Diagenode, B01020001) combined with the Bioruptor Water cooler (Cat. No. BioAcc-cool) & Single Cycle Valve (Cat. No. VB-100-0001) at high power setting (position H) using 1.5 ml TPX microtubes (Cat. No. M-50001). After centrifugation at 13 000 g for 15 min, 2.5 % of the cleared chromatin was stored at −80 °C for determining DNA input. The rest of the sample was divided in three equal parts, which were supplemented with additional 600 μl of RIPA buffer and incubated overnight at 4 °C with 14 μl of antibody-coupled protein G Dynabeads (Thermo Fisher Scientific, 10004D). Before adding the cleared chromatin, the beads were pre-blocked with bovine serum albumin and salmon sperm DNA overnight at a final concentration of 0.2 μg/μl, pre-incubated in 500 μl of RIPA buffer for 4 hours with the antibody and collected by magnetic stand to remove the unbound antibody. We used 2 μg of anti-RNA polymerase II RPB1 (Abcam, ab76123), 1 μg of anti-RNA polymerase II CTD repeat YSPTSPS (phospho S2) (Abcam, ab5095), and 3 μg of anti-CDK9 (Cell Signaling Technology, 2316S) antibodies. Additionally, 2 μg of normal mouse IgG (Santa Cruz Biotechnology, sc-2025) and normal rabbit IgG (Santa Cruz Biotechnology, sc-2027) antibodies were used to determine specificity of the signals. Validation for all antibodies is provided on the manufacturers’ websites. For the CDK9 assays, the cells were cross-linked using ChIP Cross-link Gold reagent (Diagenode, C01019027) and formaldehyde according to manufacturer’s instructions and lysed as above. Lysates were sonicated during 30 cycles of 15 seconds sonication with the Misonix XL-2000 Ultrasonic Liquid Processor using the P-1 Microprobe 3.2 mm tip, power setting 6, in 1.7 ml RNase/DNase-free microcentrifuge tubes (Sigma, CLS3620), which were kept for 1 min on ice between the cycles. Generally, 1/60th of the precipitated ChIP sample was used for each qPCR reaction. For input DNA, 1/100th of the DNA input dissolved in 100 μl of water was used for each qPCR reaction. Samples were amplified using FastStart Universal SYBR Green QPCR Master (Rox) (Sigma, 04913914001), DNA-specific primer pair, and LightCycler 480 II (Roche) machine. Primers were from Integrated DNA Technologies and designed using PrimerQuest Tool. Values were normalized to their levels in DNA inputs and calculated as a ratio to the values obtained from the DMSO control samples. Ser2-P ChIP values were further normalized to the values of total Pol II. Results from three independent experiments are presented as the mean ± s.e.m. Sequences of the primers used in ChIP-qPCR assays along with their genomic locations are listed in Supplementary Table 19.

### RNA Interference, siRNAs and antisense-oligonucleotide treatment

For RT-qPCR, cytotoxicity, viability, and apoptosis assays, cells grown on 6-well plates were transfected with 50 pmol per well of the indicated siRNA for 48 hours. For co-immunoprecipitation assays, cells grown on 10 cm plates were transfected with 400 pmol per plate of the indicated siRNA for 48 hours. For ChIP-qPCR assay, cells grown on 15 cm plates were transfected with 450 pmol per plate of the indicated siRNA for 48 hours. 7SK was depleted for 48 hours using phosphorothioate-modified antisense DNA oligonucleotide (as7SK), of which sequence was CCTTGAGAGCTTGTTTGGAGG. 100 pmol per 6-well plate well of the antisense DNA oligonucleotide was used. Transfections were performed using Lipofectamine RNAiMAX reagent (Thermo Fisher Scientific, 13778-150) according to the manufacturer’s instructions. RBM7 siRNA #1 had the sense sequence rGrCrGrUrArArArGrUrCrArGrArArUrGrArArUTT. RBM7 siRNA #2 had the sense sequence rGrGrArUrArArArGrGrCrArUrUrGrCrUrUrArATT. Control siRNA was from Qiagen (AllStars Negative Control siRNA, SI03650318). Efficiency of the knockdowns were evaluated by Western blotting or RT-qPCR assays.

### Cytotoxicity Assay

HeLa Flp-In, RPE or HFF-1 cells were seeded on 96-well plates 16 hours before experiment to ensure 80 % confluency. Concentrations of chronic treatments with 4-NQO were 250 nM (HeLa Flp-In), 500 nM (HFF-1), and 1 μM (RPE). Where indicated, 250 nM of flavopiridol was used. Doses of UV irradiation were 40 J/m^2^ (HeLa) and 60 J/m^2^ (RPE). Cytotoxicity was evaluated using CellTox Green Cytotoxicity Assay (Promega, G8741). CellTox Green Dye was added to the cells together with chemicals or immediately after UV irradiation. Fluorescence was measured at the indicated time points using PerkinElmer Victor X3 Reader. Results from three independent experiments are presented as fluorescence values relative to the untreated control and plotted as the mean ± s.e.m.

### Viability Assay

HeLa Flp-In cells were seeded on 96-well plates 16 hours before experiment to ensure 80 % confluency and subjected to the same experimental conditions as in cytotoxicity assays. Cell viability was examined using alamarBlue Cell Viability Assay (Thermo Fisher Scientific, DAL 1025). Fresh medium containing the alamarBlue cell viability reagent was added to the cells two hours prior to the indicated time points. Fluorescence was measured at the indicated time points using PerkinElmer Victor X3 Reader. Results from three independent experiments are presented as fluorescence values relative to the untreated control and plotted as the mean ± s.e.m.

### Apoptosis Assay

HeLa Flp-In cells were seeded on 96-well plates 16 hours before experiment to ensure 80 % confluency and subjected to the same experimental conditions as in cytotoxicity assays. Activation of apoptosis was assessed using RealTime-Glo Annexin V Apoptosis and Necrosis Assay (Promega, JA1011). Luminescence was measured at the indicated time points using PerkinElmer Victor X3 Reader. Results from three independent experiments are presented as luminescence values relative to the untreated control and plotted as the mean ± s.e.m.

### Statistical analysis

Differential gene expression analysis of 4sU-seq data was performed using edgeR (Robinson et al., 2010). P-values obtained from edgeR were corrected by multiple testing using the method by Benjamini and Hochberg (Benjamini and Hochberg, 1995) for adjusting the false discovery rate (FDR) and a p-value cutoff of 0.01 was applied. Data shown for all qPCR-based experiments and functional assays were collected from at least 3 biological replicates as indicated in individual figure legends and are presented as means ± s.e.m. Statistical significance and *P* values were determined by one-tailed Student’s *t* test performed between the indicated paired groups of biological replicates. Where the results were not statistically significant, *P* values are not indicated.

### Data availability

The RBM7 iCLIP data have been deposited to ArrayExpress Archive (EMBL-EBI) under the accession code E-MTAB-6475. The 4sU-seq data have been deposited to Gene Expression Omnibus (GEO) repository (NCBI) under the accession code GSE110272. Tables S3-S6 and S9 are and other data that support the findings are available upon request from the corresponding author.

## ACKNOWLEDGMENTS

We thank Céline Verheggen and Edouard Bertrand for sharing the HeLa Flp-In cell lines; Qiang Zhou for sharing anti-LARP7 antibody; Michal Lubas for providing uaRNA and eRNA annotations; Stephen P. Jackson for advice and suggestions; and Zeping Luo for technical assistance. This work was funded by Academy of Finland [1263825 and 1309846 to M.B.]; Sigrid Juselius Foundation [4704610 to M.B.]; Deutsche Forschungsgemeinschaft (DFG) [Do-1275/2-1 to L.D.]; European Research Council [206726-CLIP to J.U.]; Francis Crick Institute with core funding from Cancer Research UK (FC001002), the UK Medical Research Council (FC001002), and the Wellcome Trust (FC001002) to J.U.; Deutsche Forschungsgemeinschaft (DFG) [FR2938/7-1 to C.C.F.]; A.B. was supported in part by funding from the Doctoral Program in Biomedicine, University of Helsinki. C.R.S is supported by an Edmond Lily Safra fellowship.

## AUTHOR CONTRIBUTIONS

M.B. conceptualized the study and was assisted by A.B. and A.J.C.Q. in experimental design. M.B. supervised the study. C.R.S. performed iCLIP with the assistance from Melanie B. and analysed iCLIP data with the assistance from J.U. K.F. performed V-PAC experiments. A.B., T.H. and L.D. performed 4sU-seq, and C.C.F. analyzed 4sU-seq data. All other experiments were performed by A.B., A.J.C.Q, T.L. and P.K. M.B. wrote the manuscript with input from all co-authors.

## AUTHOR DECLARATION

The authors declare no competing interests.

